# Deep Coverage and Quantification of the Bone Proteome Provides Enhanced Opportunities for New Discoveries in Skeletal Biology and Disease

**DOI:** 10.1101/2022.11.20.517228

**Authors:** Jacob P. Rose, Charles A. Schurman, Christina D. King, Joanna Bons, Jordan B. Burton, Sandip K. Patel, Amy O’Broin, Tamara Alliston, Birgit Schilling

## Abstract

Dysregulation of cell signaling in chondrocytes and in bone cells, such as osteocytes, osteoblasts, osteoclasts, and an elevated burden of senescent cells in cartilage and bone, are implicated in osteoarthritis (OA). Mass spectrometric analyses provides a crucial molecular tool-kit to understand complex signaling relationships in age-related diseases, such as OA. Here we introduce a novel mass spectrometric workflow to promote proteomic studies of bone and cartilage. This workflow uses highly specialized steps, including extensive overnight demineralization, pulverization, and incubation for 72 h in 6 M guanidine hydrochloride and EDTA, followed by proteolytic digestion. Analysis on a high-resolution Orbitrap Eclipse and Orbitrap Exploris 480 mass spectrometer using Data-Independent Acquisition (DIA) provides deep coverage of the bone proteome, and preserves post-translational modifications, such as hydroxyproline. A spectral library-free quantification strategy, directDIA, identified and quantified over 2,000 protein groups (with ≥ 2 unique peptides) from calcium-rich bone matrices. Key components identified were proteins of the extracellular matrix (ECM), bone-specific proteins (e.g., secreted protein acidic and cysteine rich, SPARC, and bone sialoprotein 2, IBSP), and signaling proteins (e.g., transforming growth factor beta-2, TGFB2), and lysyl oxidase homolog 2 (LOXL2), an important protein in collagen crosslinking. Post-translational modifications (PTMs) were identified without the need for specific enrichment. This includes collagen hydroxyproline modifications, chemical modifications for collagen self-assembly and network formation. Multiple senescence factors were identified, such as complement component 3 (C3) protein of the complement system and many matrix metalloproteinases, that might be monitored during age-related bone disease progression. Our innovative workflow yields in-depth protein coverage and quantification strategies to discover underlying biological mechanisms of bone aging and to provide tools to monitor therapeutic interventions. These novel tools to monitor the bone proteome open novel horizons to investigate bone-specific diseases, many of which are age-related.

## Introduction

The age-related decline of skeletal health over the lifetime of an organism [1, 2] leads to bone and joint diseases, such as osteoporosis and osteoarthritis (OA). As these pathologies are tightly correlated with increasing age, their impact is more drastic in aging populations, and case rates are rising worldwide as individual lifespans increase. Indeed, global OA case rates have risen by about 10% between 1990 and 2017, and case rates in the United States of America have increased by 23% in that same time. Currently, OA prevalence in the USA is over 6,000 cases per 100,000 people as reported in 2020 [3]. Bone fragility, due to loss of bone mass, independently or together with a loss of bone quality, likewise affects a growing segment of the aging population. Unfortunately, the mechanisms by which aging propels OA or osteoporosis, like age-related neurodegeneration and other diseases, remains unclear [4], despite clinical and molecular evidence for common etiologies [4–6]. Clinically relevant assays and biomarkers for age related skeletal disease are crucially needed to diversify diagnostic and treatment options and to monitor treatment efficacy.

Skeletal biology is a complex and multi-faceted ecosystem of cell types, including bone-building osteoblasts and bone-resorbing osteoclasts. Osteocytes, embedded within the calcified bone matrix, represent 90–95% of all cells within the bone. These cells signal and communicate with osteoblasts and osteoclasts directly via dendritic processes and through secretion of proteins that can influence their formation and activity Through mechanisms that remain unclear, osteocytes support the homeostasis of chondrocytes that reside within and maintain articular cartilage. Aging and increased cellular senescence burden impact all cell types within bone and cartilage, alter their homeostatic function, and lead to tissue degeneration (**Figure 1)** [2, 7–9]. Interestingly and encouragingly, interventions that target and reduce senescent cell burden in mice prevent age-related bone loss in mice [10].

**Figure 1.**
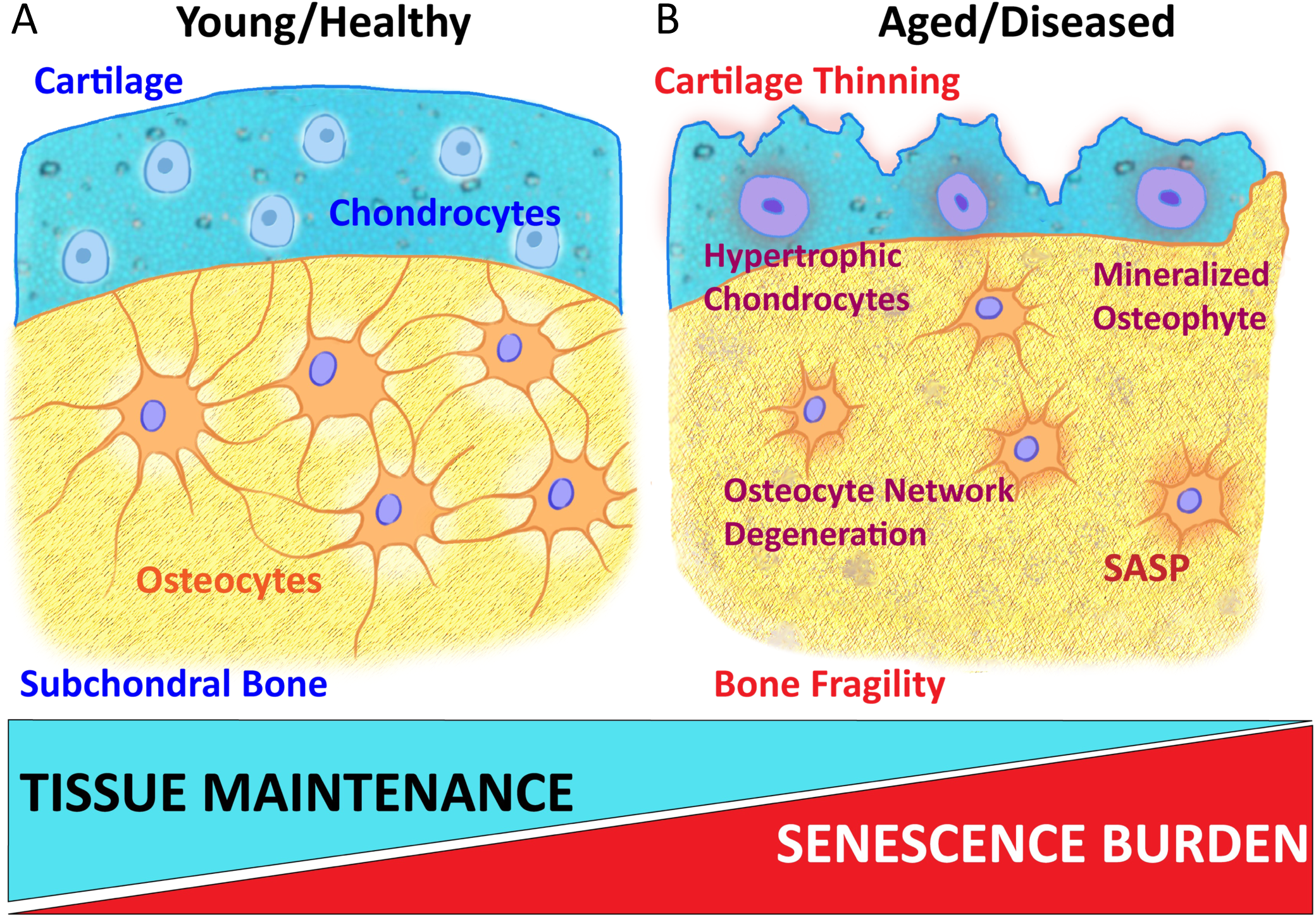
Comparison of bone microenvironments in healthy and aged/diseased states. As bones age and the senescence burden increases, the bone microenvironment changes, which is implicated in bone fragility and in osteoarthritis. Additionally, osteocyte lacunocanalicular networks lose canaliculi, and chondrocyte homeostasis is disrupted, causing cartilage degeneration that is accompanied by chondrocyte hypertrophy and osteophyte (bony spurs) formation.

With age, the balance between osteoblast regulated bone deposition and osteoclast mediated bone resorption is disrupted: bone resorption begins to outpace bone deposition, leading to lost bone mass [2, 11, 12]. Osteocyte dendritic processes typically decrease significantly with age [13, 14], and other mechanisms that maintain bone tissue material properties decline [15–19]. Thus, aging in the skeleton is associated with both osteopenia and osteoporosis, clinical conditions of low bone mass, that lead to increased rates of fracture in aged-populations, as well as fragility fractures in individuals with clinically normal bone mass [20, 21]. Additionally, joint disease progression, such as OA, is associated with advancing age as the cellular and material regulatory mechanisms maintaining cartilage decline [22, 23]. Temporal alterations to skeletal cell behavior, including a shift of homeostatic mechanisms towards more inflammatory phenotypes, also contribute to the breakdown of skeletal tissues and lead to age-related diseases.

One cellular mechanism associated with both inflammation and aging is cellular senescence. During senescence, homeostatic cells exit their normal cell cycles and adopt a senescence-associated secretory phenotype (SASP) that is characterized by a host of inflammatory secreted factors [24–26]. The role that the SASP and senescence play in bone health is well documented [27, 28], but also understudied on a molecular level. Recent efforts examined biomarkers of senescence and senescence burden in multiple disease pathologies, including OA [29, 30]. Skeletal tissue is especially susceptible to damage via the SASP as secreted factors may not only alter skeletal cell behavior but directly damage skeletal tissues that enable locomotion [27, 31]. While the SASP contains a broad range of factors, several of the most well identified markers include matrix metalloproteinases, other proteolytic enzymes, and ATPase ion pumps that regulate intracellular and pericellular pH [25, 32]. These factors are normally well regulated by skeletal cells, and are used to maintain tissue quality, but with age their unchecked action can directly damage skeletal tissues leading to age-related skeletal disease. In addition, many of the intricate and well-regulated molecular signaling pathways that maintain balanced bone deposition and resorption can be members of, or targeted by, SASP. For instance, both sclerostin (SOST) and receptor activator of NF-κβ ligand (RANKL) are secreted proteins produced by osteocytes that increase in serum with age. These factors are known to be induced during cellular senescence within bone cells and directly shift bone remodeling towards bone resorption by suppressing osteoblast bone formation and increasing osteoclastic bone resorption, respectively [10, 33–36]. Dysregulation of signaling in skeletal cells and an elevated senescent burden are implicated in bone aging and in progressive loss of mechanical function that contributes to bone frailty, osteoporosis, and osteoarthritis [1, 9]. With the important roles of secreted proteins in both skeletal tissue remodeling and managing skeletal cell type function, along with their altered behavior in age and senescence, there is a dire need to apply molecular tools that directly measure protein presence and quantitative regulation in the context of age-related skeletal diseases.

Large-scale unbiased analytical techniques, such as transcriptomics, proteomics, metabolomics, and lipidomics, are powerful methods that are often used to gain comprehensive insights into complex biological systems and their changes in aging and diseases. Several successful transcriptomics and single-cell-sequencing efforts have been undertaken in bones [37–39], but few quantitative bone proteomic studies have been published [40–45]. Given the structural relevance of extracellular matrix (ECM) proteins and signaling peptides within the skeleton, analyzing bone proteome profiles during aging and age-related skeletal biology will be highly beneficial and insightful. Proteomic studies appear to be limited in skeletal tissues, possibly due to the complex, dense and mineralized matrix, and overall analytical challenges of efficient protein extraction using existing protocols. The mineralized ECM of bone contains large amounts of calcium in the form of matrix-bound hydroxyapatite, Ca_10_(PO_4_)_6_(OH)_2,_ that is embedded within and around collagen fibrils. These hydroxyapatite minerals cause nonspecific interactions between positively charged amino acids with phosphate groups and carboxyl residue complexes with calcium [46], and thus, conventional protein lysis extractions often fail or are inefficient. Protein extraction techniques that disrupt nonspecific interactions and carboxyl residue complexes between proteins and hydroxyapatite are critically needed for efficient and comprehensive protein coverage in bone.

The protein compositions within bone and other skeletal tissues are highly specialized in form and function. Assessment of the skeletal proteome would give insight into the dominant constituent, collagen, as well as non-collagenous ECM proteins and cellular proteins. Uniquely, post-translation modifications of individual pro-collagen molecules through proline oxidation are required to stabilize the triple-helix structure of collagen itself through stereo-electric effects or water-bridged hydrogen bonding [47]. Genetic disorders, such as osteogenesis imperfecta [48] and others that interrupt hydroxyproline modifications are phenotypically characterized by brittle bones in young patients [49] and resemble some age-related pathologies. Capturing post-translational modifications (PTMs) and their interactions among structural proteins within the skeleton is critical for assessing bone health, and proteomic methods that preserve these alterations are crucial for studying skeletal tissues.

With this study, we present a novel proteomic workflow for an unbiased analysis of skeletal tissues to better understand the mechanisms underlying osteoarthritis, osteoporosis, and other age-related skeletal pathologies by examining the changes within the proteomic landscape between bone and cartilage. Deep protein coverage and quantification are achieved with our novel robust and reproducible proteomics protocol to efficiently extract proteins from the complex bone matrix and the subsequent combination with modern quantitative mass spectrometric strategies. We adapted and improved steps from a reported method that extracted proteins from bones [50], and that was also recently used by our group for analysis of cystinuric bladder stones [51]. To investigate bone aging and diseases we paired this efficient extraction protocol with data-independent acquisition mass spectrometry (DIA-MS) [52–54], and this novel workflow profiles and accurately quantifies both dynamic changes in protein composition, as well as changes in protein PTMs. The application of quantitative proteomics to further understand bone cell signaling dynamics in the context of aging and cellular senescence is expected to lead to the discovery of novel biomarkers and to discover new therapeutic targets for the treatments of age-related bone diseases.

## Results

### Reproducible protein extraction from mouse bones and proteomic analysis

Our novel workflow comprises highly-specialized steps, including collection of long bones from C57/BL6 mice, removing and flushing the bone marrow, followed by extensive demineralization overnight using 1.2 M HCl, pulverization with a bead mill homogenizer, incubation for 72 h in 6 M guanidine hydrochloride and EDTA to extract proteins, a buffer exchange to remove the guanidine hydrochloride and, finally, proteolytic digestion of the extracted protein lysates (see **Figure 2**). Protein concentrations were initially determined with a bicinchoninic acid assay (BCA), as a first step of quality control and to demonstrate successful protein extraction. Subsequently, bone protein lysates were prepared for integrated chromatographic and mass spectrometric (MS) analysis, including tandem mass spectrometry (MS/MS).

**Figure 2.**
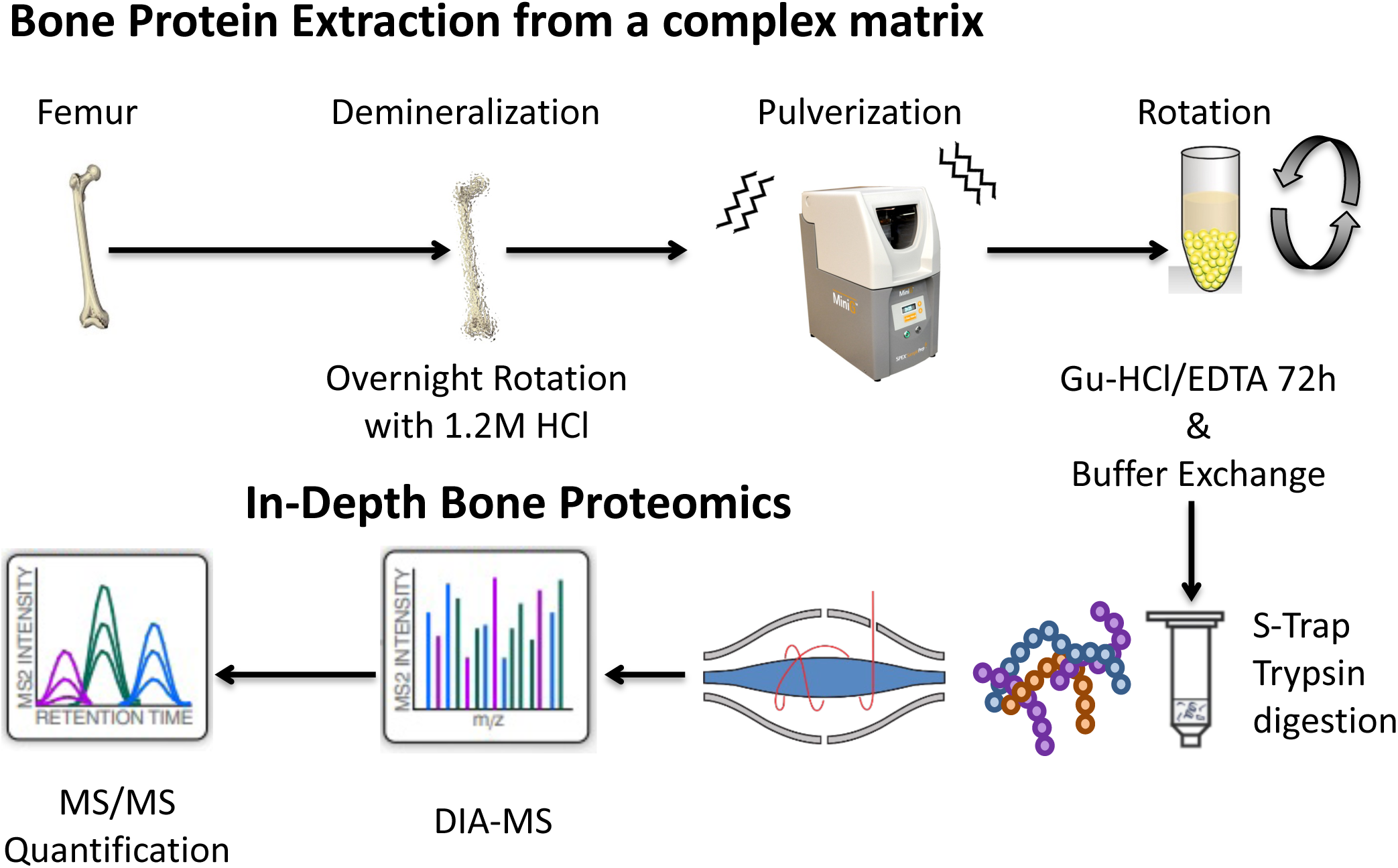
Sample workflow and strategy for Mass Spectrometry analysis. Femurs and tibia from wild-type mice were demineralized by rotating overnight in 1.2 M HCl and pulverized with a SPEX bead mill at 1500 strokes per minute for 2 minutes, the samples were allowed to cool in liquid nitrogen and pulverized for 2 more minutes as above. Proteins were extracted for 72 hours at 4° C in 6 M guanidine HCl. The extracted protein lysates were buffer exchanged into Tris-HCl to remove guanidine and digested with Strap columns and trypsin overnight. Subsequently, the proteolytic digestions were analyzed by MS in DIA mode for an in-depth proteome analysis on either the Orbitrap Exploris 480 or the Orbitrap Eclipse Mass Spectrometer.

In order to assess the performances of our workflow and reduce biological variability in the initial MS experiment, we pooled extracted protein lysates from 2 tibia and 2 femurs from a single wild type mouse. We then proteolytically digested with trypsin, and peptides were analyzed on an Orbitrap Exploris 480 in five technical replicates. Briefly, we used a label-free, highly quantitative DIA-MS approach, in which the sequential selection of MS1 precursor ion windows (or m/z segments) for MS/MS enable that the entire MS1 mass range was subjected to fragmentation within each scan cycle throughout the entire gradient [52, 54]. This comprehensive acquisition utilized a precursor ion isolation scheme that consisted of 26 variable windows and covered an *m/z* range of 350–1,650 with an overlap of 1 *m/z* per each DIA window [55]. Data files were processed using a spectral library-free strategy, referred to as directDIA within the Spectronaut algorithm (Biognosys) [56, 57].

Acquisitions were assessed for quality by displaying their retention time regressions thereby plotting the indexed retention time in relation to the observed retention time with a representative graph shown in **Figure 3a**. Each point along the curve represents a measured precursor ion and its retention regression. These indexed retention time regressions were normalized for slight variations in measured retention times from acquisition to acquisition, and thus improve the quantification accuracy. DIA-MS acquisitions assayed reproducibility by measuring coefficients of variation (CV). **Figure 3b** visualizes the precursor ion CV distribution correlating the precursor ion CV values against the respective measured peak area abundances and indicating high reproducibility. As expected, very low abundant analytes appear to be more variable. However, across five MS replicates, a median CV of 8.7% was reported for precursor ions, and 85% of precursor ions featured a CV that was smaller than 20% (**Figure 3c-d and Supplemental Table S1**).

**Figure 3.**
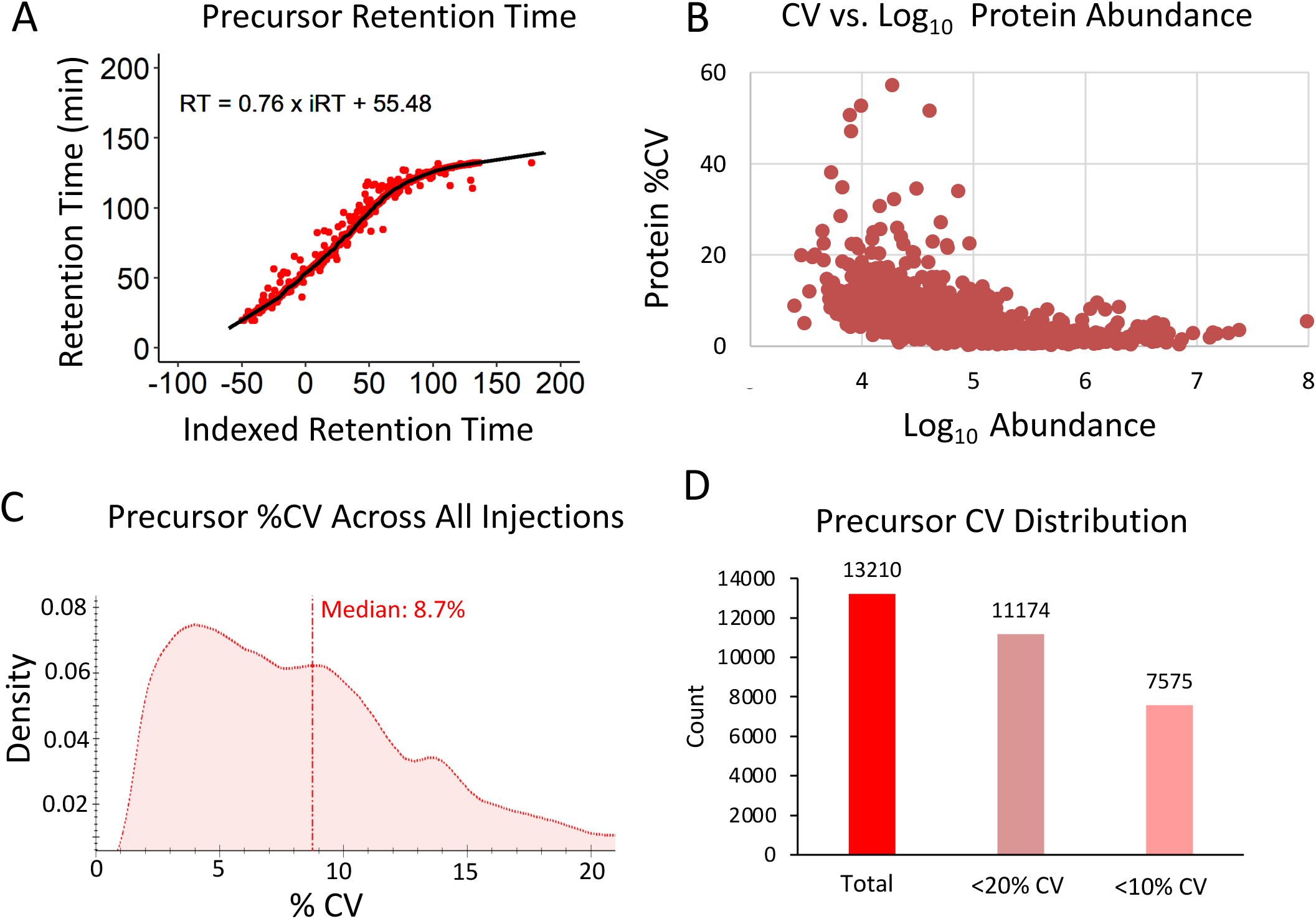
Quality assessment of pooled bone sample workup injected in five technical replicates. Data acquisitions from the 5 technical replicates of the pooled sample were assessed using A) precursor ion retention time (RT) regressions to illustrate acquisition to acquisition reproducibility and RT normalization, B) coefficient of variation (CV) vs abundance to determine precursor ion confidence, C) precursor ion %CV density to visualize precursor ion variability, and D) precursor ion CV distribution to quantify precursor ion reproducibility.

### In-depth Coverage of Cellular and Extracellular Matrix (ECM) Proteins from Mouse Femurs

In a separate study, a cohort of five mice was utilized to isolate individual femurs, followed by our optimized workflow for protein extraction and proteolytic digestion. Mass spectrometric DIA analysis of the digested protein lysates on an Orbitrap Eclipse (Thermo) from this independent cohort of mice resulted in the confident identification and quantification of 2,108 protein groups, identified with ≥2 unique peptides **(Supplemental Table S2)**.

We performed various Gene Ontology (GO) analyses to determine which cellular compartments and biological processes were enriched in this bone-derived protein dataset. GO cellular compartments, such as the cytoplasm, nucleus, plasma membrane, and the extracellular region, presented most of the annotated identifications and showed that a variety of proteins were included in this proteome analysis (**Figure 4a**). In addition, a biological process GO analysis showed broad enrichment for diverse cellular processes: both intracellular and extracellular processes (e.g., oxidative phosphorylation, tricarboxylic acid cycle, and collagen metabolic processes) and processes related to the mechanical function of bone (e.g., collagen fibril organization, collagen catabolism, endochondral bone growth, and musculoskeletal movement) as shown in **Figure 4b and Supplemental Table S3.** These results highlight a major strength of the sample preparation process, because capturing diverse protein profiles that include highly abundant ECM components, such as collagens and proteoglycans, and metabolic or signaling proteins, such as TGF-beta ligands, is needed for a mechanistic understanding of bone biology. To more completely interrogate bone-specific proteins, we assessed protein coverage and pseudo-MS/MS spectrum quality for secreted protein acidic and cysteine rich (SPARC), a protein typically secreted by osteoblasts during bone formation [58]. We observed a 51% sequence coverage for SPARC, with the observed tryptic peptides displayed in green **(Figure 4c)**. In addition, we confirmed the extracted ion chromatogram (XIC) quality, reproducibility, and intensity for several y-fragment ions in replicates WT1 and WT3 **(Figure 4d)**.

**Figure 4.**
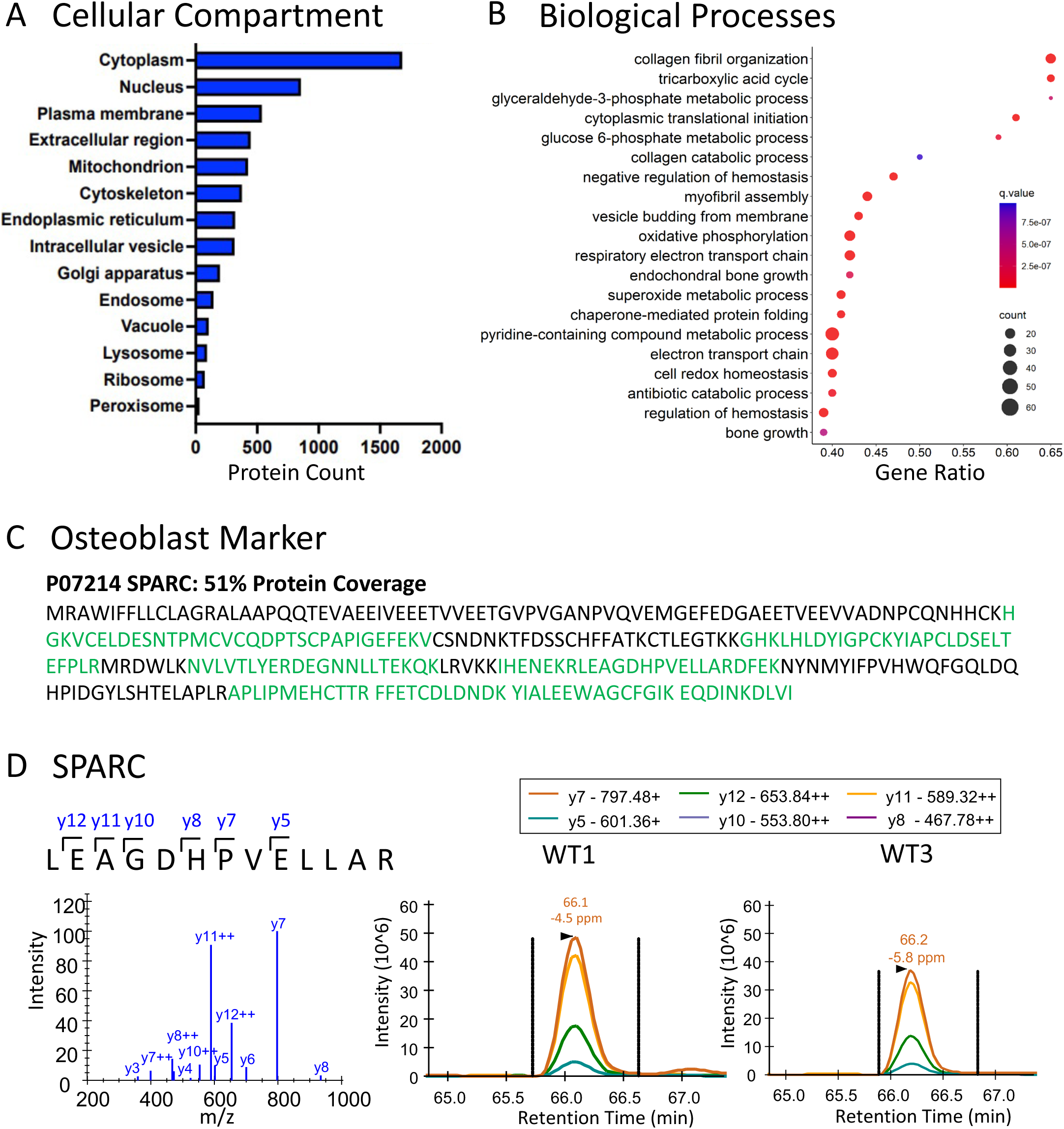
Proteomic results and Gene Ontology analysis for five wild-type femurs. Proteins identified in wild-type mouse femurs were analyzed using ConsensusPathDB-mouse (Release MM11, 14.10.2021) GO for both A) “cellular compartment” and B) “biological processes.” The most enriched cellular compartments were the cytoplasm and nucleus while the most significant GO Biological Process was “collagen fibril organization”. C) Protein sequence coverage for secreted protein acidic and cysteine rich (SPARC), an osteoblast marker, is highlighted where green sections of the sequence (51%) are confidently identified in the proteomics data. D) directDIA generated Pseudo-MS/MS spectrum (constructed from DIA generated spectral library) and extracted ion chromatogram of the SPARC peptide LEAGDHPVELLAR (m/z=710.38, z=2+) illustrates peak quality and intensity for 2 selected samples, WT1 and WT3.

### Detection of Functionally Diverse Collagens and Formation of Structural Networks

We next examined the coverage of collagens identified using our novel bone proteomics methodology. Collagens are a diverse family of proteins that are important for bone biology because of the large structural formations that are formed from many interacting collagens [59] [60]. In our analysis, we identified major functional types of collagens denoted by color and labeled A-F. This included fibrillar collagens (e.g., collagens type I, II, III, and V) (**Figure 5a**) which accounted for over 78% of total collagen abundance as displayed in the vertical bar graph on the right. The next most abundant family of collagen identified was the Beaded Filament collagens, mostly made up of distinct chains of Collagen VI (**Figure 5e**). The third most abundant functional family identified, made up mostly of Collagen XII, was the “fibril-associated collagens with interrupted triple helices”, or FACIT family (**Figure 5b** ∼10% of total collagens) including collagen types IX, XII, XIV, XVI, and XXII. Interestingly, we identified multiple members of the thrombospondin family (Thbs1-4), including Thbs1 which is the major protein involved in structural assembly and FACIT biology [61]. Other families of collagens were also identified in smaller amounts including Hexagonal Network Collagens, (**Figure 5c**) Network Collagens, (**Figure 5d**) and other, anchoring collagens **(Figure 5f).**

**Figure 5.**
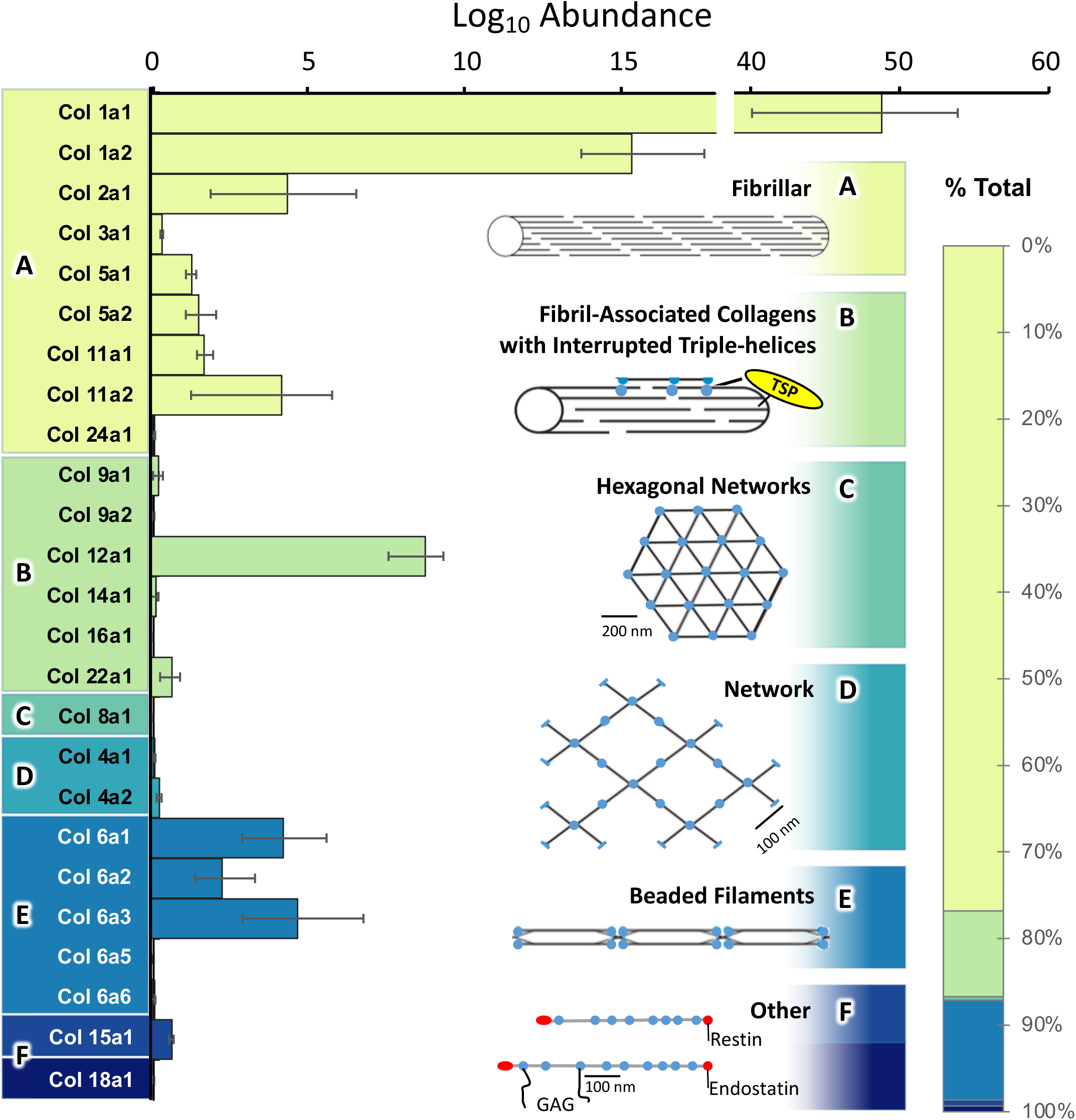
Overview of collagen subtypes identified in five wild-type mouse femurs. A horizontal bar graph shows the estimated abundance of individual collagen proteins confidently identified in the analysis, for example, fibrillar collagens account for 78% of total collagen abundance, mostly from Collagen type 1. Additionally, collagen structural families are quantified and denoted in panels A-F with their percent contribution of total collagen abundance as represented in the vertical bar graph on the right. A: Fibrillar, B: FACIT’s, C: Hexagonal Networks, D: Networks, E: Beaded Filaments, F: Other.

### Preservation of Hydroxyproline Modifications

PTMs are biologically relevant for many tissue types, [62] in bone, we analyzed hydroxyproline modifications, including the determination of proline PTM site-localization within the measured peptides and proteins. Hydroxyproline is an important PTM, specifically in collagens, that helps stabilize 3-dimensional protein-protein interactions among different collagen chains [47, 59]. To confidently identify and site-localize hydroxyproline modifications (+ 15.99 Da), we initially processed our samples using data-dependent acquisition (DDA), where precursor ions are isolated in ‘tight’ windows (1 m/z), and are then subjected to MS/MS, providing less complex and highly specific MS/MS spectra. To generate true quantitative assays and to query these modifications using quantitative DIA acquisitions, we generated a spectral library that included the 1,382 DDA-identified hydroxyproline-containing peptides using Spectronaut. Importantly, we determined that hydroxyproline modifications are preserved during our sample preparation, and can be subjected to these new quantitative assays. This search resulted in 1,186 hydroxyproline modified peptides with 85% site localization probability, corresponding to 75 modified proteins **(Supplemental Table S4B)**. In fact, our novel MS workflow allowed the identification of a hydroxyproline modified residue Pro-707 (in bold below), a previously reported [63] and structurally relevant site on the collagen alpha-1(I) chain. The modified peptide is shown (GDTGAPoxGA**Pox**GSQGAPoxGLQGMPoxGER, Col1a1), featuring both the MS/MS and the extracted ion chromatogram (XIC) **(Figure 6a).** In this example we confirmed the presence of 4 hydroxyproline residues with direct and indirect ion evidence including the y_4_, y_5_, y_10_, y_16_, and y_19_ ions. In addition, we determined that Pro-533 (GLTGS**Pox**GSPGPDGK, Col1a1) was modified and displayed a nearly complete MS/MS spectrum that provided accurate mapping of the hydroxy modification using differentiating ions like y_4_, y_6_, and y_9_ **(Figure 6b)**. These results exhibit the power of this analysis to confidently capture and localize biologically relevant PTMs without prior sample enrichment that is commonly required for PTM studies.

**Figure 6.**
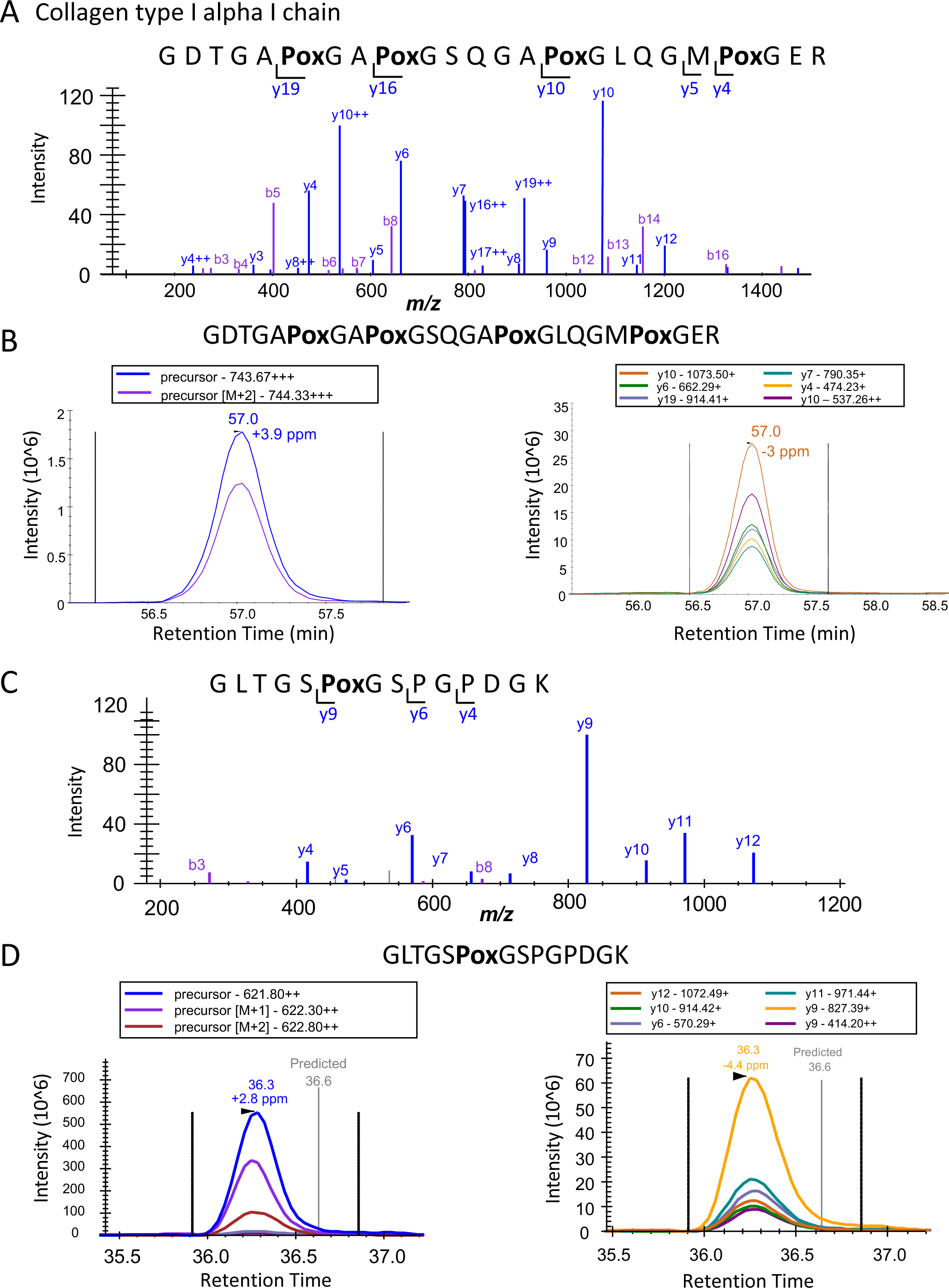
Analysis of hydroxyproline modifications on collagen I alpha chain I. Two tryptic peptides of collagen 1 alpha chain 1 (COL1A1) with highly-confident hydroxyproline site localizations due to specific and comprehensive fragmentation. A) Representative spectrum displaying the ion series for GDTGA^7**04**^**Pox**GA^7**07**^**Pox**GSQGA^7**13**^**Pox**GLQGM^7**19**^**Pox**GER. Fragmentation series provides site localization evidence to confirm modified prolines based on direct and indirect ion evidence including the y_4_, y_5_, y_10_, y_16_, and y_19_ ions. B) Extracted ion chromatograms (XIC) of GDTGA^7**04**^**Pox**GA^7**07**^**Pox**GSQGA^7**13**^**Pox**GLQGM^7**19**^**Pox**GER precursor ion (m/z=743.67, z=3+) on the left and fragment ions (y_10_, y_4_, and y_19_) on the right. C) Similar representative spectrum displaying the fragment ions for the singly modified GLTGS**^533^Pox**GSPGPDGK. Direct ion evidence for the modified peptide can be attributed to the y_4_, y_6_, and y_9_ ions. D) XIC of GLTGS **^533^Pox**GSPGPDGK precursor ion (m/z=621.80 and z= 2+) on the left and fragment ions (y_9_, y_6_, and y_10_) on the right.

### Quantification of biologically relevant ECM components and senescence markers

This dataset was initially compared to the core matrisome published by Naba et al. [64–66] to determine the level of coverage, compared to this rich proteomic dataset. We found that 42% of bone proteins identified overlapped with the known murine core matrisome (**Supplemental Table S5a**), suggesting that extracted bone proteins are largely composed of ECM and ECM-related proteins. We compared our dataset to each component of the core matrisome, such as glycoproteins, collagens, proteoglycans, ECM affiliated proteins, ECM regulator proteins, and secreted proteins **(Figure 7a)**. Specifically, we identified 70 glycoproteins and 71 ECM regulators that are part of the core matrisome. In addition to the matrisome data base, we analyzed our results in reference to a proteomic analysis of the Senescence Associated Secretory Phenotype (SASP) published by Basisty et al. [25]. In the latter publication our group outlined commonly identified proteins in the SASP using multiple inducers of senescence, considering them the Core SASP. The results from this comparison showed that 131 Core SASP factors are identified in this analysis. These common identifications include ECM modifying proteins, such as matrix metalloproteinases (MMP2), lysyl oxidase-like 2, biglycan, TIMP metallopeptidase inhibitor 2 (TIMP2), and serpin H1. Additionally, insulin signaling proteins, including multiple insulin-like growth factors, and certain members of the immune system, such as macrophage migration inhibitory factor (MIF) and high mobility group box 1 (HMGB1), were identified **(Figure 7b)**. Indeed, many of the Core SASP proteins have been quantified here, and the comparisons presented describe the relevance and depth of this proteomic workflow.

**Figure 7.**
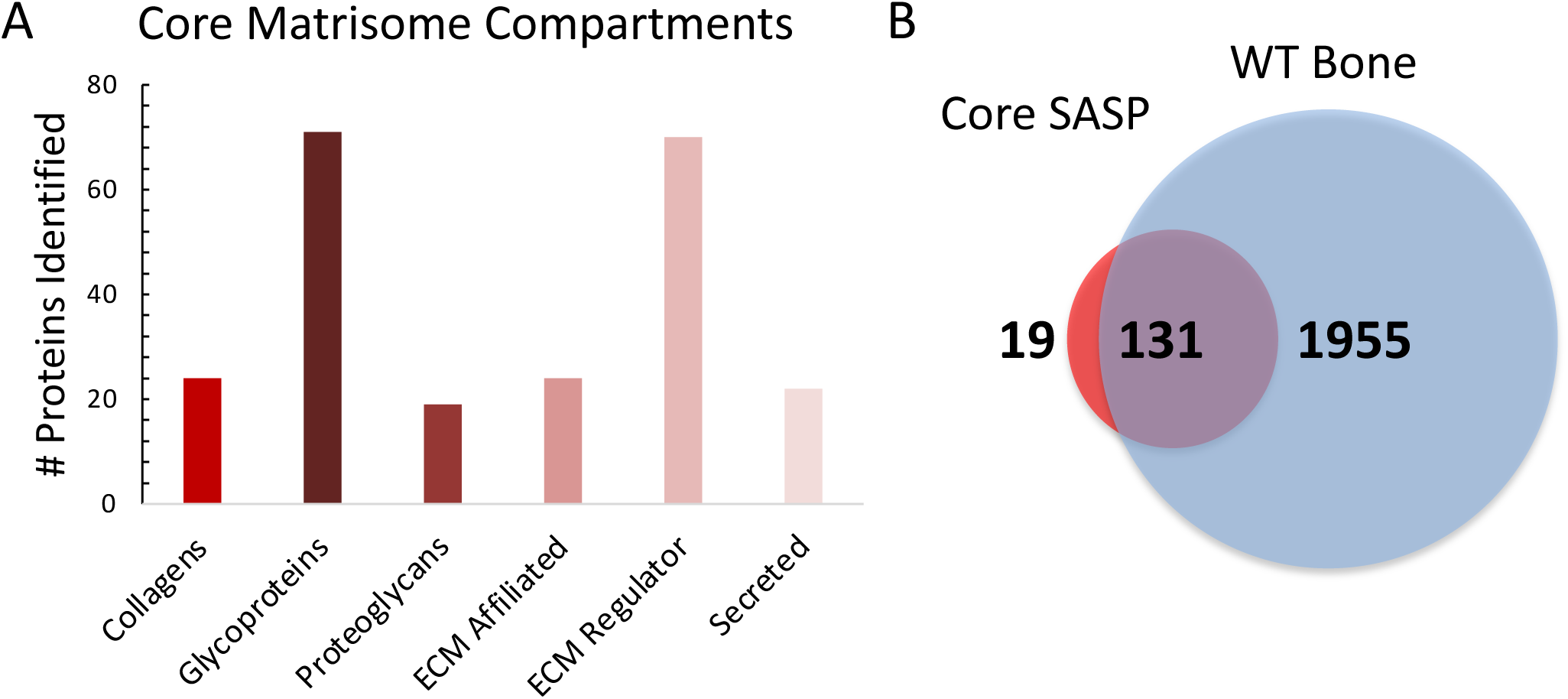
Comparison of five wild-type mouse femurs to the core matrisome database and the Core SASP. A) Proteins identified from wild-type mouse femurs were compared to components of the core matrisome showing most representation to least representation as follows: Collagens (24 proteins), glycoproteins (71 proteins), proteoglycans (19 proteins), ECM affiliated proteins (24 proteins), ECM regulators (70 proteins), secreted proteins (22 proteins) [64–66]. B) Additional comparisons of the bone proteome to the Core SASP show 131 overlapping protein identifications and only 19 unique core SASP proteins [25].

## Discussion

A novel demineralization and extraction protocol was developed to obtain deep proteomic profiles for bones to study age-related bone conditions, such as OA. Proteins were extracted and digested, and proteolytic peptides were analyzed using DIA-MS for identification and quantification of relevant molecular signatures in bone. Many bone-specific proteins and proteins important to skeletal biology were identified, including important ECM proteins, collagen-specific PTMs that influence collagen structural assembly, and potential senescence/SASP factors. Additionally, proteins identifying specific bone cell types, such as osteopontin/bone sialoprotein 1 (Opn/Bsp-1) for osteoblasts, matrix extracellular phosphoglycoprotein (Mepe) for osteocytes, and tartrate-resistant acid phosphatase type 5(TRAP) for osteoclasts, were found and confirmed bone cellular identity in our analysis **(Supplemental Figure 1)**. This proteomic protocol was optimized to overcome challenges in the bone matrix that interfere with accurate protein identification and quantification. The initial demineralization step was crucial and greatly helped to preprocess the lysates by removing calcium phosphate deposits while leaving ECM and other proteins intact in the organic lysate that remained. The other protocol steps were designed to provide an extremely efficient and reproducible workflow using S-trap columns for purification and digestion of the samples and HLB cartridges for desalting of the proteolytic peptides.

The samples were then acquired in DIA mode and processed using directDIA (Spectronaut). Searching the acquisitions in this spectral library-free fashion saves sample and instrument acquisition time. This resulted in a comprehensive bone proteome profile featuring tissue-specific, and potentially disease-relevant protein candidates. To enhance our study at the post-translational level, we investigated bone samples with DDA, and identified PTM-containing proteolytic peptides. Subsequently, we built custom PTM spectral libraries and were able to use these to extract hydroxyproline containing peptides from our DIA data, especially focusing on collagen I. Additionally, senescence markers relevant to OA progression, such as metalloproteinase 2 (MMP2), metalloproteinase inhibitors 1 and 2 (TIMP1 and 2), and signaling proteins, were also identified. Identification of well-characterized SASP factors, including HMGB1, illustrated the potential to use proteomics to assay bone senescence. Indeed, the effects of cellular senescence in other related tissues, such as cartilage, have been implicated but not characterized with rigorous proteomics, especially after proposed treatments for OA, senolytics, or other cartilage rejuvenation therapeutics. The ECM greatly influences cell signaling and cell function and is a significant determinant of cell fate by sensing mechanical stimuli from the stroma [67–69].

Since bone tissue is robustly composed of ECM, changes in the ECM proteome will be at the forefront of understanding bone biology and mechanics. These results will provide insight to other tissues where the ECM or mechano-sensing pathways have also been shown to be critically important, such as in the muscle [70], cartilage [71, 72], breast[73], ovary [74], as well as multiple cancers associated with these tissues [65, 75]. Proteomics studies are uniquely positioned to capture relevant information about collagen ratios, senescence markers, and disease contributors in the bone microenvironment using a highly confident and quantitative mass spectrometric analysis.

This new integrated proteomic workflow combining the thorough bone protein extraction protocol with a comprehensive DIA-MS analysis will allow us to analyze human and other mammalian bone samples, specifically in the context of bone fragility and osteoarthritis. We will be able to dynamically monitor disease progression, and potentially discover new biomarkers and provide tools to monitor therapeutic interventions.

## Materials and Methods

### Reagents and Standards

HPLC solvents (e.g., acetonitrile and water) were obtained from Burdick & Jackson (Muskegon, MI). Reagents for protein chemistry (e.g., iodoacetamide, dithiothreitol, guanidine hydrochloride, EDTA, and formic acid) were purchased from Sigma Aldrich (St. Louis, MO). Proteomics grade trypsin was from Promega (Madison WI). HLB Oasis SPE cartridges were purchased from Waters (Milford, MA).

### Animal Tissues

Femurs and tibia from 16 week old male mice on a C57BL/6 background were collected and cleaned of muscle and other soft tissue and marrow removed via centrifugation. Animals were housed and collected in accordance with the Institutional Animal Care and Use Committee of the University of California San Francisco. Over their lifetime, animals were housed in a pathogen-free environment with the temperature maintained between 68 °F and 74 °F, humidity between 30% and 70%, with a 12-h light/dark cycle and access ad. libitum to water and rodent chow (LabDiet 5053).

### Protein Extraction from Bones

The femurs from 5 C57/B6J (WT) male mice at age 16 weeks were stripped of any muscle, epiphyses were cut off, and the marrow was removed via centrifugation to obtain ‘clean bone tissue’ and to reduce contribution from the other tissues to the bone analysis. Because the proteomic preparation and MS analysis occurred a few days after bone harvest and bone isolation, the bones were briefly stored at -80 °C after being wrapped in Hank’s Balanced Salt Solution (HBSS) soaked gauze. Bones were demineralized by adding 1 mL of 1.2 M HCl and rotating them overnight at 4 °C [50]. The demineralized bones were subsequently transferred to a new Eppendorf tube and kept on dry ice until pulverization. All components of the homogenizer, including sample tubes and homogenizing plates, were cooled with liquid nitrogen. Individual frozen femurs from 5 mice were each pulverized with the SPEX SamplePrep 1600 MiniG tissue homogenizer in polycarbonate tubes with a 9.5-mm steel grinding ball for 2 minutes at 1500 strokes/minute. After the first 2 minutes, the samples were removed from the homogenizer, allowed to cool in liquid nitrogen for 3 minutes, and homogenized for another 2 minutes at 1500 strokes/minute. Pulverized samples were transferred into a fresh Eppendorf tube using 800 µL of extraction buffer (6 M guanidine hydrochloride, 10 mM Tris-HCl, 50 mM EDTA) and incubated by rotating at 4 °C for 72 hours [76]. Subsequently, the samples were spun for 3 minutes at 15,000 x *g* to separate bone matrix and the supernatants. Supernatants containing the soluble proteins were buffer exchanged to remove guanidine hydrochloride with Amicon 3 kDa Centrifugal Filters. Samples were spun through the filter at 12,000 x *g* for 20 minutes and resuspended in 500 µL of 10 mM Tris-HCl (pH 7). This wash was spun down at 12,000 x *g* for 20 minutes, and the procedure was repeated 2 more times for a total of three washes with 10 mM Tris-HCl. The final addition of 10 mM Tris-HCl was spun as described above until the samples were reduced to 20 µL. Extracted proteins in 10 mM Tris-HCl were quantified using a bicinchoninic acid assay (BCA). During our protocol optimization steps, we were able to achieve efficient protein yields of 50-80 µg (assessed by BCA) from a single mouse femur, that would typically have an original wet weight of ∼50 mg.

### Proteolytic Digestion

For each individual bone sample, 20 µg of protein lysate was brought up to 4% SDS using a 10% SDS solution. Samples were then reduced using 20 mM dithiothreitol in 50 mM triethylammonium bicarbonate buffer (TEAB, pH 7) at 50 °C for 10 minutes, cooled to room temperature (RT) and held at RT for 10 minutes, and alkylated using 40 mM iodoacetamide in 50 mM TEAB (pH 7) at RT in the dark for 30 minutes. Samples were acidified with 12% phosphoric acid to obtain a final concentration of 1.2% phosphoric acid. S-Trap buffer (90% methanol in 100 mM TEAB at pH ∼7.1) was added, and samples were loaded onto the S-Trap mini spin columns (Protifi, Farmingdale, NY). The entire sample volume was spun through the S-Trap mini spin columns at 4,000 x *g* at RT, binding the proteins to the mini spin columns. Subsequently, S-Trap mini spin columns were washed twice with S-Trap buffer at 4,000 x *g* at RT and placed into clean elution tubes. Samples were initially digested for 1 hour at 47°C with sequencing grade trypsin (Promega, San Luis Obispo, CA) dissolved in 50 mM TEAB (pH 7) at a 1:25 (w:w) enzyme:protein ratio. Finally, an additional trypsin solution was added at the same w:w ratio, and proteins were digested overnight at 37°C to ensure more complete peptide cleavage. Peptides were sequentially eluted from the mini S-Trap spin columns with 80 µL of 50 mM TEAB, 0.5% formic acid (FA) in water, and 50% acetonitrile (ACN) in 0.5% FA. After centrifugal evaporation, samples were resuspended in 0.2% FA in water and desalted with Oasis 10 mg Sorbent Cartridges (Waters, Milford, MA). The desalted elutions were subjected to centrifugal evaporation, and they were re-suspended in 0.2% FA in water at a final concentration of 1 µg/µL. Finally, indexed Retention Time Standards (iRT, Biognosys, Schlieren, Switzerland) were added to each sample, according to manufacturer’s instructions [77].

### Sample Pooling Strategy

The initial proteomic experiment (**Figure 3**) featured one pooled sample with lysates from 2 tibia and 2 femurs from a single mouse, prepared as described for the individual femurs above, injected in 5 technical replicates on the Orbitrap Exploris 480 (**Supplemental Methods 1**). This experiment only used long bones from a single mouse to reduce biological variability and to ensure a deep coverage of the bone proteome. The next experiment (**Figure 4**) featured femurs from 5 different mice to assess biological variability, but were only from one bone type.

### Mass Spectrometric Analysis using Data-Independent Acquisition (DIA) and Data-Dependent Acquisition (DDA)

Reverse-phase HPLC-MS/MS analyses were performed in DIA mode on a Dionex UltiMate 3000 system coupled online to an Orbitrap Eclipse Tribrid (Thermo Fisher Scientific, San Jose, CA). The solvent system consisted of 2% ACN, 0.1% FA in water (solvent A) and 98% ACN, 0.1% FA in water (solvent B). The following text describes the chromatography schema for the MS acquisitions, flowrates will be described as “volume of solvent/ 1 minute”. For the DIA acquisitions, digested peptides (200 ng) were loaded onto an Acclaim PepMap 100 C_18_ trap column (0.1 x 20 mm, 5-µm particle size; Thermo Fisher Scientific) over 5 min at 5 µL/min with 100% solvent A. Peptides were eluted on to an Acclaim PepMap 100 C_18_ analytical column (75 µm x 50 cm, 3-µm particle size; Thermo Fisher Scientific) at a flow rate of 300 nL/min using the following gradient (indicating the % of solvent B): 2% B for 5 min, linear from 2% to 20% B in 95 min, linear from 20% to 32% B in 20 min, increase to 80% B in 1 min, hold at 80% B for 9 min, and back to 2% B in 1 min. The column was re-equilibrated for 29 min with 2% of solvent B/98% solvent A, and the total gradient length was 160 min. Each of the samples was acquired in DIA mode [52, 54, 55] in technical duplicates and in DDA mode. For DIA, survey MS1 spectra were collected at 120,000 resolution (Automatic Gain Control (AGC) target: 3e6 ions, maximum injection time: 60 ms, 350–1,650 *m/z*), and MS2 spectra at 30,000 resolution (AGC target: 3e6 ions, maximum injection time: Auto, Normalized Collision Energy (NCE): 27, fixed first mass 200 *m/z*). The DIA isolation scheme consisted of 26 variable windows covering the 350–1,650 *m/z* range with an overlap of 1 *m/z* per each window **(Supplemental Table S6)**. For the DDA acquisitions, digested peptides (200 ng) were loaded onto an Acclaim PepMap 100 C_18_ trap column (0.1 x 20 mm, 3-µm particle size; Thermo Fisher Scientific) over 10 min at 2 µL/min with 100% solvent A. Peptides were eluted on to an Acclaim PepMap 100 C_18_ analytical column (75 µm x 50 cm, 3-µm particle size; Thermo Fisher Scientific) at 300 nL/min using the following gradient (indicating the % of solvent B): 2% B for 10 min, linear from 2% to 20% B in 95 minutes, linear from 20% to 32% B in 20 min, increase to 80% B in 1 minute, hold at 80% B for 9 min, and back to 2% B in 1 min. The column was re-equilibrated for 29 min with 2% of solvent B/98% solvent A, and the total gradient length was 165 min. For DDA, survey MS1 spectra were collected at 240,000 resolution (AGC target: 1.2e6 ions, maximum injection time: Auto, 350–1,500 m/z). Precursor ions with a charge state 2–5+ and an intensity above 2e4 were automatically selected for HCD fragmentation at NCE 27 in the orbitrap for a cycle time of 3 s. MS2 spectra were collected at 30,000 resolution (AGC target: 1e5 ions, maximum injection time: Auto, fixed first mass 200 *m/z*). Dynamic exclusion was set to 60 s.

### Data-Independent Acquisition (DIA) Data Processing

DIA data files were processed in Spectronaut v16 (Biognosys) using directDIA. Data were searched against the *Mus musculus* reference proteome with 58,430 entries (UniProtKB-TrEMBL), accessed on 01/31/2018. Data extraction parameters were set as dynamic, and non-linear iRT calibration with precision iRT was selected. Trypsin/P was set as the digestion enzyme, and two missed cleavages were allowed. Cysteine carbamidomethylation was set as a fixed modification, and methionine oxidation and protein N-terminus acetylation were set as dynamic modifications. For the protein level, identification was performed requiring a 1% q-value cutoff on the precursor ion and protein levels. Unique protein groups were reported with at least two unique peptide identifications. The protein level quantification was based on the peak areas of extracted ion chromatograms (XICs) of 3–6 MS2 fragment ions, specifically b- and y-ions, with local normalization and q-value sparse data filtering applied. In addition, iRT profiling was selected.

### DDA Spectral Library Generation and DIA Quantification for Hydroxyproline-containing Peptide Level Analysis

A DDA spectral library was generated in Spectronaut v16 using slightly modified Biognosys default settings (BGS), and the same *Mus musculus* database. Briefly, for the Spectronaut Pulsar search, trypsin/P was set as the digestion enzyme, and two missed cleavages were allowed. Cysteine carbamidomethylation was set as fixed modification, and proline oxidation, methionine oxidation, and protein N-terminus acetylation were set as variable modifications. Identifications were validated using 1% false discovery rate (FDR) at the peptide spectrum match (PSM), peptide and protein levels, and finally the best 3–6 fragment ions per peptide were kept. The spectral library contains 20,705 peptides and 2,372 protein groups, including 1,382 hydroxyproline-containing peptides corresponding to 85 hydroxyproline-containing protein groups (**Supplemental Table S7**). Identification was performed requiring a 1% q-value cutoff on the precursor ion and protein levels. The PTM site localization score was selected with a probability cutoff of 0.75. For PTM analysis, DIA data were processed in Spectronaut v16, using the generated DDA spectral library (described above). Quantification was based on extracted ion chromatograms (XICs) of 3 – 6 MS2 fragment ions, specifically b- and y-ions, without normalization and data filtering using q-value sparse. Grouping and quantitation of PTM peptides were accomplished using the following criteria: minor grouping by modified sequence and minor group quantity by mean precursor ion quantity.

#### Pathway Analysis

Over-representation analysis was performed using Consensus Path DB-mouse (Release MM11, 14.10.2021), developed by the bioinformatics group at the Max Planck Institute for Molecular Genetics (Berlin, Germany) [78, 79]. The list of quantifiable proteins was used to evaluate which gene ontology terms, including biological processes, molecular functions, and cellular components, were significantly enriched in these samples. Gene ontology terms identified from the over-representation analysis were subjected to the following filters: q-value < 1.0e^-6^, term category = b (biological processes), and term level ≥ 3. Dot plots were generated using the ggplot2 package [80] in R (version 4.0.5; RStudio, version 1.4.1106) to visualize significantly enriched biological processes from each comparison **(Supplemental Table S3)**.

## Acknowledgement/Funding

We acknowledge the support from the National Institutes of Health NIH: NIA (U01 AG060906, PI: Schilling, and T32 AG000266 to Schurman; PI: Campisi/Ellerby), the Office of the Director for the Orbitrap Eclipse system (1S10 OD028654-01, PI: Schilling), NIDCR (R01 DE019284, PI: Alliston). In addition, we are grateful to the Glenn Foundation (to Patel), and the forever healthy Foundation (PI: Schilling).

---

**Supplemental Figure 1.**
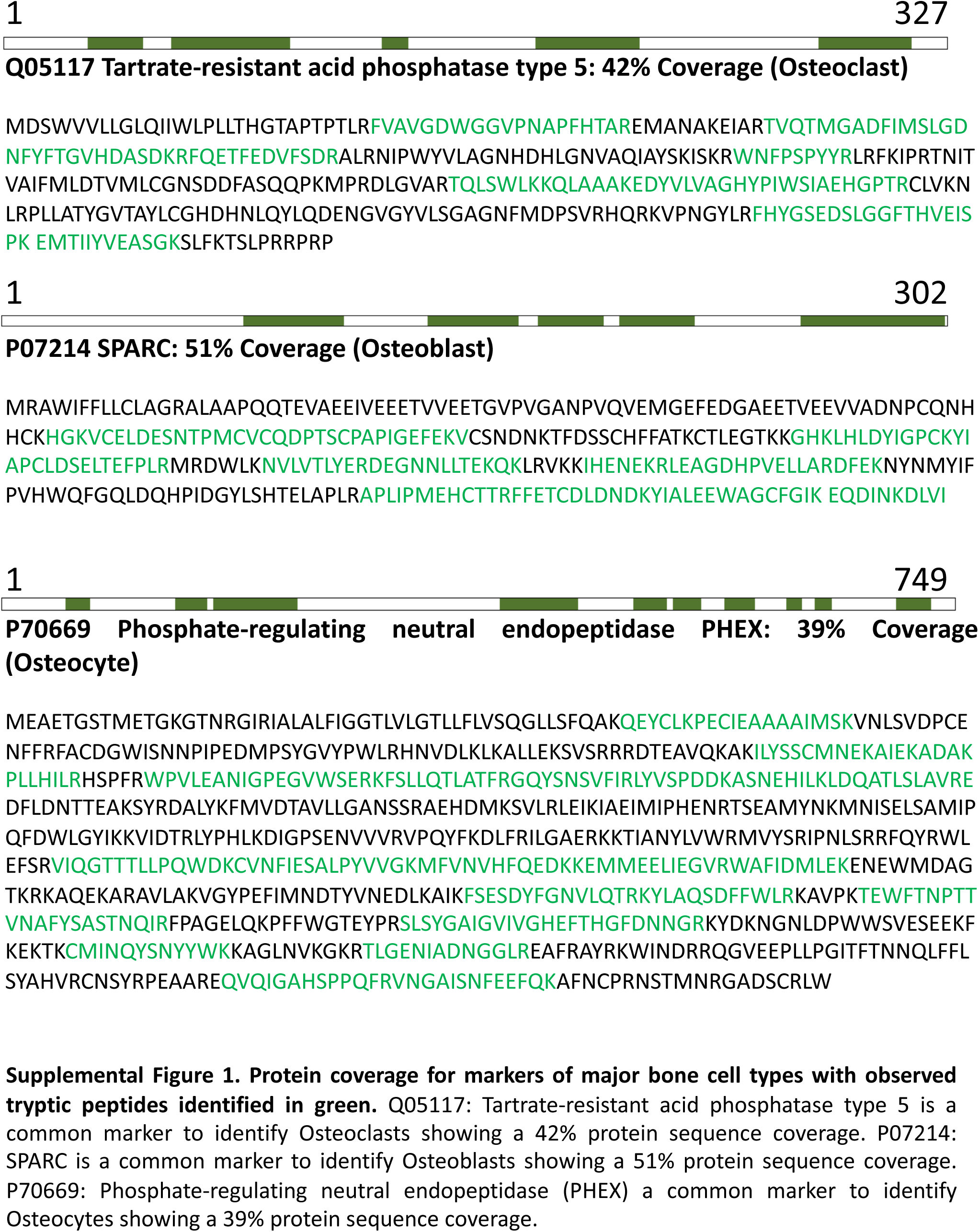
Protein coverage for markers of major bone cell types with tryptic peptides in the sequence shown in green. Q05117: Tartrate-resistant acid phosphatase type 5 is a common marker to identify Osteoclasts showing a 42% peptide coverage. P07214: SPARC is a common marker to identify Osteoblasts showing a 51% peptide coverage. P70669: Phosphate- regulating neutral endopeptidase (PHEX) a common marker to identify Osteocytes showing a 39% peptide coverage.

